# Elraglusib induces cytotoxicity via direct microtubule destabilisation independently of GSK3 inhibition

**DOI:** 10.1101/2024.07.06.602326

**Authors:** Josh T. Coats, Shuyu Li, Tomoyuki U. Tanaka, Sudhir Tauro, Calum Sutherland, Adrian T. Saurin

**Affiliations:** Division of Cellular & Systems Medicine, School of Medicine, University of Dundee, Dundee, UK; Division of Molecular Cell and Developmental Biology, School of Life Sciences, University of Dundee, Dundee, UK; Division of Molecular & Clinical Medicine, School of Medicine, University of Dundee, Dundee, UK

## Abstract

Elraglusib (9-ING-41) is an ATP-competitive inhibitor of glycogen synthase kinase-3 (GSK3) with pre-clinical studies demonstrating broad activity against many tumour types. Promising early-phase clinical trial data led to FDA orphan drug status, and a randomized phase 2 study in combination with cytotoxic chemotherapy in pancreatic cancer has recently completed its recruitment. Similarly, single-agent responses in adult T-cell leukaemia/lymphoma and melanoma, and combination treatment data in several other tumour types have been encouraging. The elraglusib mechanism of action is unknown, but it is unlikely to act through GSK3 inhibition because cytotoxicity is observed below the IC_50_ for GSK3 and other small molecule GSK3 inhibitors do not produce cytotoxic effects, at least in lymphoma cells. We show here that elraglusib perturbs chromosomal alignment to cause a mitotic arrest in multiple tumour lines. This arrest is caused by direct microtubule depolymerisation, which prevents the attachment of kinetochores to microtubules. At clinically relevant doses, these mitotically arrested cells eventually undergo mitotic slippage, leading to gross chromosome missegregation, DNA damage and apoptosis. These effects explain the cytotoxicity of elraglusib because temporarily pausing cell cycle progression with the CDK4/6 inhibitor palbociclib abolishes any drug-induced genotoxicity and apoptosis. In summary, elraglusib acts as a potent direct microtubule destabilizer both *in vitro* and across multiple cancer types, resulting in mitotic arrest, DNA damage and apoptosis. These effects likely account for its broad pan-cancer activity, which does not rely upon GSK3 inhibition as they are not replicated by other GSK3 inhibitors.

## Introduction

Glycogen synthase kinase-3 (GSK3) is a ubiquitously expressed serine/threonine kinase critical for many cellular processes, including proliferation and apoptosis (1-3). The two GSK3 paralogs, α and β, are expressed from independent genes and are 98% homologous within the catalytic domain (4). Higher GSK3 expression has been associated with poorer outcomes in some cancers, and it has, therefore, been proposed as a potential therapeutic target (5, 6).

Elraglusib (9-ING-41), developed as an ATP-competitive small molecule GSK3β inhibitor, has demonstrated pre-clinical efficacy against a wide variety of tumour types, including lymphoma, pancreas, ovarian, bladder, renal, breast, and glioblastoma (6-13). Early evidence of clinical efficacy led to FDA orphan drug status for pancreatic cancer, and a randomized phase 2 trial in combination with gemcitabine and nab-paclitaxel has recently completed recruitment (14). Phase 1 and case-study data with elraglusib monotherapy also suggest potential efficacy in adult T-cell leukaemia/lymphoma and melanoma (15, 16).

We have previously shown that the anti-lymphoma effects of elraglusib occur below the GSK3α/β IC_50_, cannot be replicated with other structurally distinct small molecule GSK3 inhibitors and are unaffected by *GSK3A/B* knockdown, thereby questioning the role of GSK3 in its mechanism of action (17, 18). We set out to establish the mechanism of action of elraglusib since this will enhance clinical trial design and may improve the development of its anti-cancer efficacy.

## Results

### Elraglusib induces a mitotic arrest in lymphoma cells

Elraglusib has been reported to induce cell cycle arrest at G2 and/or M-phases in cancer cell lines based upon cellular DNA content analysis using propidium iodide (6, 9, 10). Therefore, we sought to investigate these effects further. To distinguish between a G2 (4N) and M-phase (mitotic 4N) arrest, we performed flow cytometry using propidium iodide and anti-phospho-Histone H3 Ser10 (H3-pS10) staining. Elraglusib treatment for 24 hours in Karpas-299 (K299) lymphoma cells, led to a dose-dependent accumulation of H3-pS10 positive mitotic cells, and, to a lesser extent, 4N cells that are H3-pS10 negative (Fig. 1A, B). Statistically significant changes were observed at elraglusib concentrations at or above 1µM. These are clinically relevant doses because phase 1 trial data shows elraglusib reaches peak plasma concentration around 17-20µM, and remains above 1µM for 24 hours post-infusion (15). Similar experiments with two structurally distinct GSK3 inhibitors, CT99021 and LY2090314, produced no cell cycle arrest in either cell line (Fig. 1C, D) at concentrations shown to inhibit GSK3 activity to a greater extent than that achieved by elraglusib (17). We conclude that elraglusib primarily causes a mitotic arrest and that these effects are not due to inhibition of GSK3.

**Figure 1.**
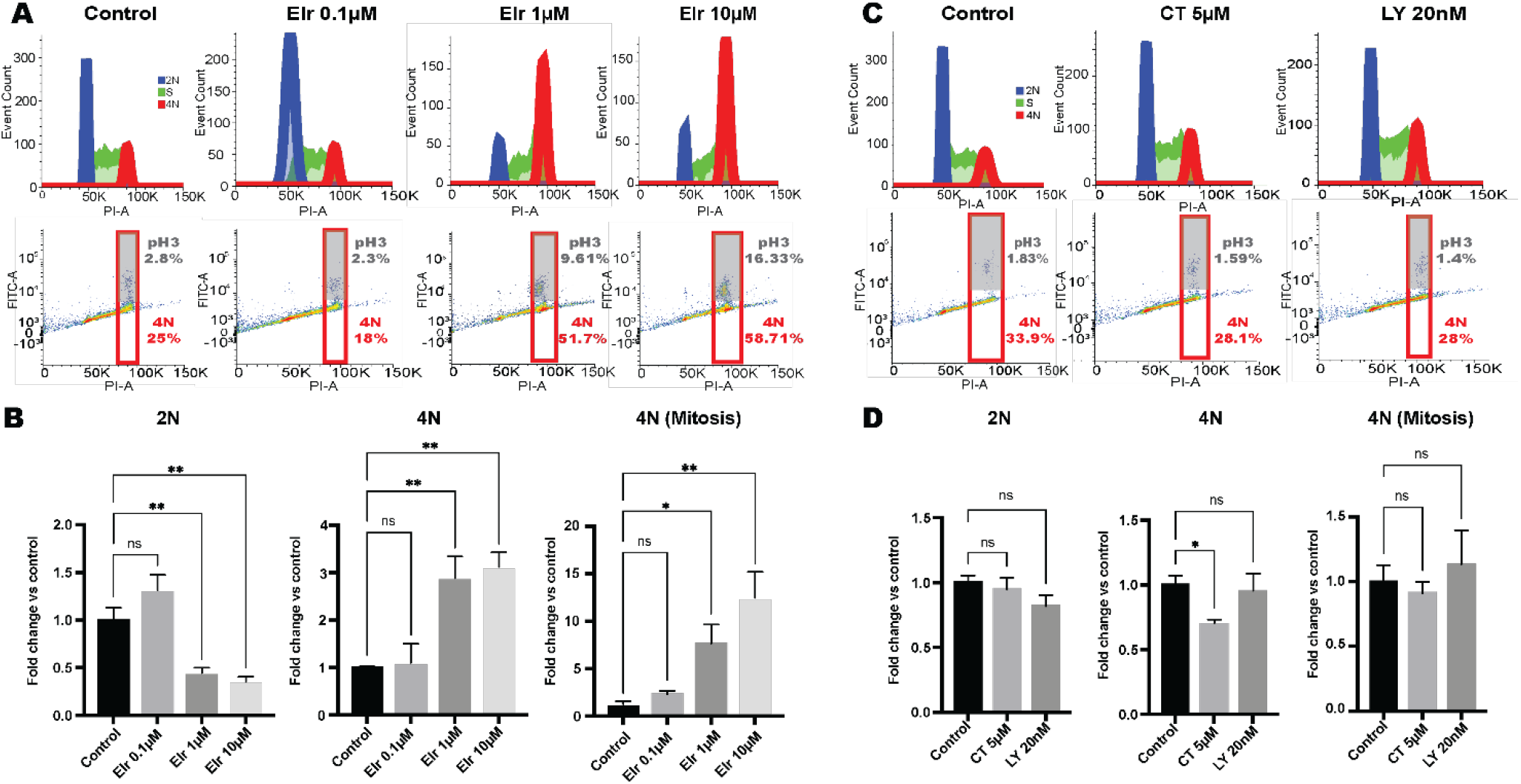
Elraglusib causes a mitotic arrest. **(A-B)** Example flow cytometry plots (A) and quantifications (B) showing the effect of 24h elraglusib (Elr) treatment on the percentage of 2N, 4N and 4N-mitotic cells. **(C-D)** Example flow cytometry plots (C) and quantifications (D) following 24h treatments with two structurally distinct GSK3 inhibitors, CT99021 and LY2090314. Karpas-299 cells were treated for 24 hours prior to cell cycle analysis by flow cytometry using propidium iodide and anti-phospho-histone-H3 (ser10) staining. 10,000 events were analysed per sample. Cell cycle histograms are representative of 3 independent experiments, and quantifications combine 3 independent experiments. Fold change calculated versus control, error bars are SEM, statistical significance by one-way ANOVA p ≤0.05.

### Elraglusib impairs kinetochore-microtubule attachment to cause a mitotic arrest

To understand why elraglusib causes a mitotic arrest, we performed live cell imaging of U2OS cells expressing H2B-GFP. Whilst control cells or GSK3-inhibited cells completed mitosis within a typical 30-minute timeframe, cells treated with elraglusib (at ≥1µM) were delayed in mitosis for many hours (Fig. 2A). This mitotic delay was associated with a failure to align chromosomes correctly (Fig. 2B and supplementary movies 1 and 2). Cells remained arrested in this state until the end of the movie, or they exited mitosis with unaligned chromosomes, in a process known as mitotic slippage (Fig. 2B, C). Mitotic slippage is frequently observed following microtubule-based therapies, leading to apoptosis or senescence in G1 (19).

**Figure 2.**
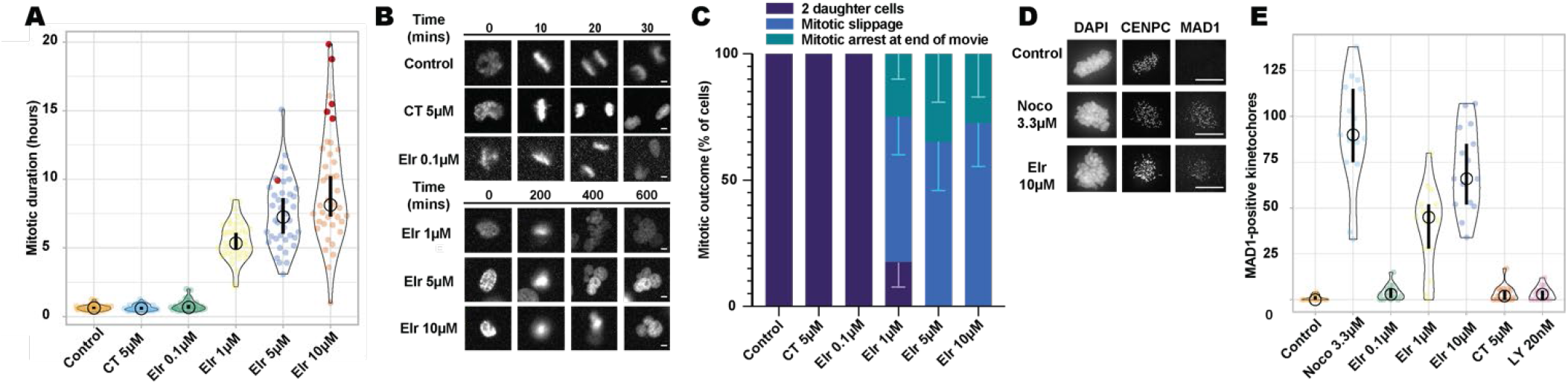
Elraglusib impairs kinetochore-microtubule attachment to cause mitotic arrest and mitotic slippage (A-C). Mitotic duration (A) with CT99021 (CT) and elraglusib (Elr), example images (B), and mitotic fate quantifications (C). U2OS H2B-GFP/mCherry-α-tubulin cells were treated and imaged every 5 minutes for 24 hours. 10 cells were analysed per field of view, with two fields per condition for each experiment. Mitotic duration (A) and fate plots (C) combine two independent experiments. For graph A, red data points indicate the movie ended with the cell remaining in mitosis, and the thick vertical lines represent a 95% confidence interval (CI) around the median, which can be used for statistical comparison of treatments by eye (see methods). For graph C, error bars represent SD. **(D, E)**. Example images (D), number of MAD1-positive kinetochores with nocodazole (Noco), elraglusib (Elr), CT99021 (CT) and LY2090314 (LY) (E). MCF7 cells were synchronized in G1 with palbociclib for 24 hours, then released for 16 hours in complete media before a 2-hour RO3306 block, wash off and MG132 treatment for 30 mins. Cells were treated for 10 mins with nocodazole (Noco), elraglusib (Elr), CT99021 (CT) or LY209014 (LY) before fixing and immunofluorescence microscopy.

CENPC was used to identify kinetochores, and MAD1 positivity was used to assess kinetochore attachment. 5 mitotic cells were imaged per coverslip. Images are representative of 3 independent experiments. Quantification represents MAD1-positive kinetochores from 5 mitotic cells per condition and combines 3 independent experiments. For graph E, the thick vertical lines represent a 95% confidence interval (CI) around the median, which can be used for statistical comparison of treatments by eye (see methods). Scale bars = 10μM We hypothesized that these defects were due to an inability to properly attach chromosomes to microtubules via the kinetochore. In this situation, the spindle assembly checkpoint (SAC) is activated to delay mitotic exit (20). This is due to the presence of MAD1 on unattached kinetochore, which generates the SAC signal until kinetochores becomes attached to microtubules and MAD1 is removed. Immunofluorescence imaging demonstrated that 10 minutes of elraglusib treatment significantly increased the number of unattached, MAD1 positive, kinetochores in MCF7 cells (Fig. 2D, E). Fully depolymerising microtubules with 3.3µM nocodazole (21) also increased MAD1 kinetochore occupancy to near 100%, as expected; however, GSK3 inhibition with CT99021 or LY2090314 had no effect. We conclude that elraglusib delays chromosome alignment by inhibiting kinetochore-microtubule attachment, leading to a persistent mitotic arrest that eventually results in mitotic slippage. These effects cannot be explained by inhibition of GSK3.

### Elraglusib acts by directly destabilising microtubules

Kinetochore-microtubule attachment errors could be due to changes at the kinetochore and/or the microtubules. To assess the latter, we performed immunofluorescence imaging of mitotic MCF7 cells treated for 24 hours with elraglusib (Fig. 3A). This demonstrated defects in mitotic spindle assembly, similar to the effects observed with low doses of nocodazole, implying microtubule polymerisation could be inhibited. To probe this further, we performed tubulin fractionation experiments to directly quantify the proportion of polymerized and depolymerized microtubules, which appear in the insoluble and soluble fractions, respectively (22). Figures 3B and C demonstrate that treatment of MCF7 cells with elraglusib for 5h caused a dose-dependent decrease in insoluble tubulin and a corresponding increase in soluble tubulin. This was similar to the effects observed with the microtubule depolymerizer nocodazole, but opposite to those observed with the microtubule stabilizer paclitaxel (Fig. 3B, C). Similar effects were also observed in K299 (Fig. 3D, E) and HH cells (Fig. 3F, G).

**Figure 3.**
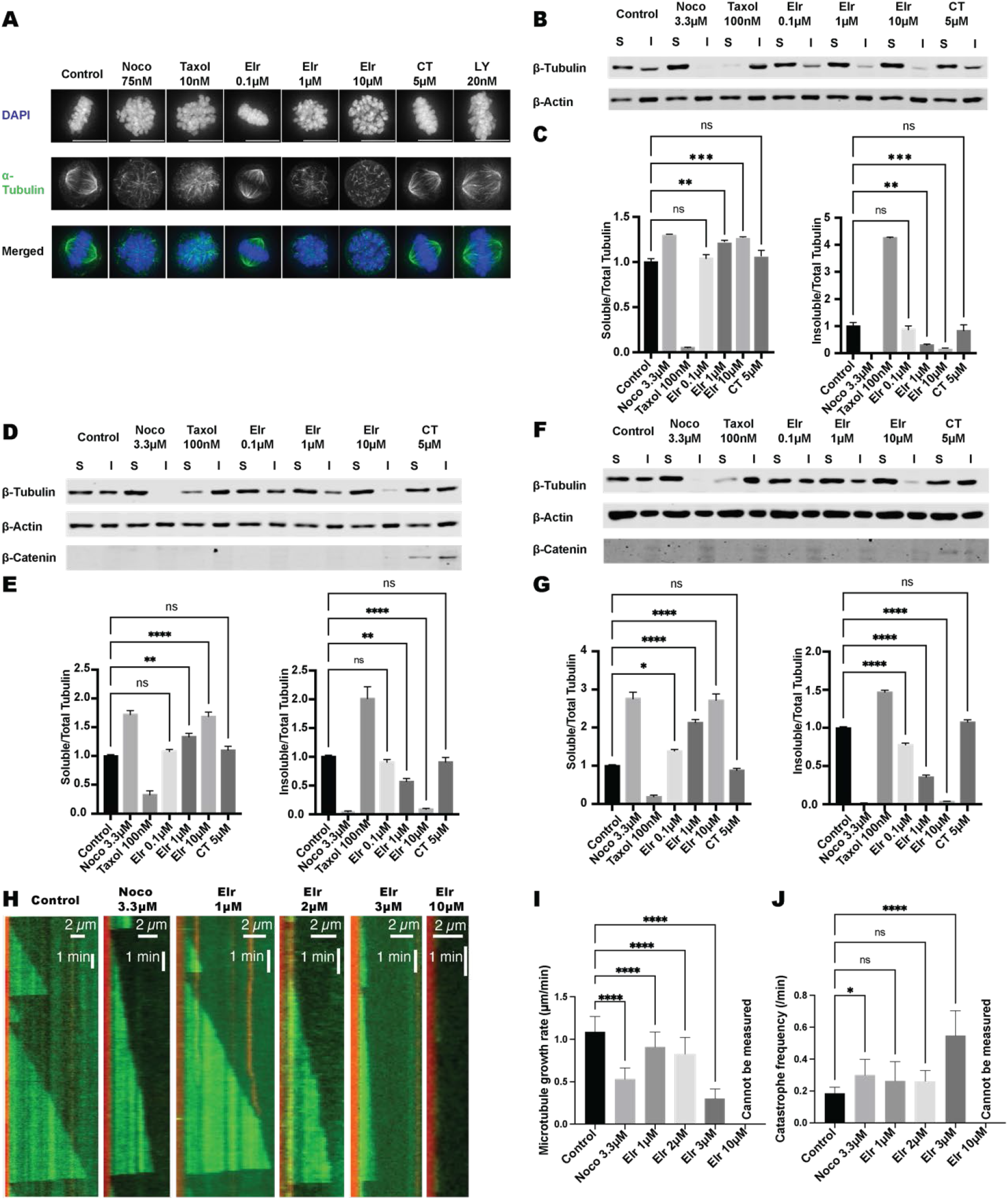
Elraglusib acts via direct microtubule destabilisation. **(A-J)**. Example images **(A)** of MCF7 cells treated with nocodazole (Noco), paclitaxel (Taxol), elraglusib (Elr) and CT99021 (CT). MCF7 cells were treated for 24 hours before fixation and immunofluorescence microscopy. 10 mitotic cells were imaged per coverslip, images are representative of 3 independent experiments. Tubulin fractionation experiments comparing soluble (S) and insoluble (I) tubulin by immunoblot in MCF7 **(B, C)**, Karpas-299 **(D, E)** and HH **(F, G)** cells. Cells were treated for 5 hours and lysed in microtubule stabilisation buffer before fractionation into soluble (S) and insoluble (I) tubulin and analysis by immunoblot. Quantification was performed by normalising to actin loading control and then to the experimental control. Blots are representative of 3 independent experiments, and quantification combines 3 independent experiments. Fold change calculated versus control, error bars are SEM, statistical significance by one-way ANOVA p ≤0.05. Scale bars = 10μM. **(H)** Kymographs illustrating dynamics of microtubule growth *in vitro* in the absence of drug (control) or the presence of 3.3µM nocodazole, 1, 2, 3 or 10µM elraglusib. Rhodamine-labelled short stable microtubules were seeded on the coverslip, and 12µM tubulins (containing 4% fluorescent HiLyte 488 tubulins) were added to the reaction to generate and visualize dynamic microtubules. Scale bars = 2µM **(I, J)** Quantification of microtubule growth rate (I) and catastrophe frequency (J) with no drug treatment (control) or with 3.3 µM nocodazole, 1, 2, 3 or 10 µM elraglusib. The number of analysed microtubule growth events: n = 34 for control; n =42 for 3.3 µM nocodazole; and n = 43, 24, and 21 for 1, 2, and 3 µM elraglusib, respectively. Microtubule catastrophe frequency was analysed for dynamic microtubules on individual microtubule seeds – the number of analysed microtubule seeds: n=13 for control; n=20 for 3.3 µM nocodazole; and n=13, 18, and 15 for 1, 2, and 3 µM elraglusib, respectively. Error bars are SD. Statistical significance by one-way ANOVA p≤0.05.

In almost all cases, elraglusib affects microtubule solubility at or above 1µM, which is consistent with the concentrations needed to cause mitotic delays and unattached kinetochores. We also observed no effect on microtubule solubility with the GSK3 inhibitor CT99021 at 5µM, despite the fact that the downstream GSK3 target β-catenin was stabilized at this concentration, indicating significant GSK3 inhibition (2) (Fig. 3B-G).

To test whether elraglusib directly induces microtubule destabilisation independently of microtubule-associated proteins or motor proteins, we performed *in vitro* microtubule growth assays (Fig. 3H-J). Elraglusib impaired microtubule growth in a dose-dependent manner (Fig. 3I) and increased the frequency of microtubule catastrophe (Fig. 3J). These effects were similar to those observed with nocodazole. No clear microtubules were observed at the highest concentration of elraglusib used (10µM), suggesting significant tubulin depolymerisation.

### Elraglusib-induced genotoxicity and apoptosis are abolished by a cell cycle arrest

Mitotic slippage commonly leads to DNA damage and apoptosis (19). We hypothesized that this could explain the cytotoxicity observed following elraglusib treatment. In agreement, elraglusib treatment led to a significant increase in γH2AX and induced PARP-cleavage in both K299 (Fig. 4A, B) and HH cells (Fig. 4C, D), compared to DMSO-treated controls. These levels were similar to or higher than observed following treatment with the DNA-damaging agent doxorubicin. Importantly, pre-treatment with palbociclib 1µM for 24 hours to arrest cells in G1 prior to elraglusib treatment essentially abolished these effects (23). This demonstrates that the genotoxic effects of elraglusib require cell cycle progression, most likely because cells must transit through mitosis and undergo mitotic slippage to promote apoptosis.

**Figure 4.**
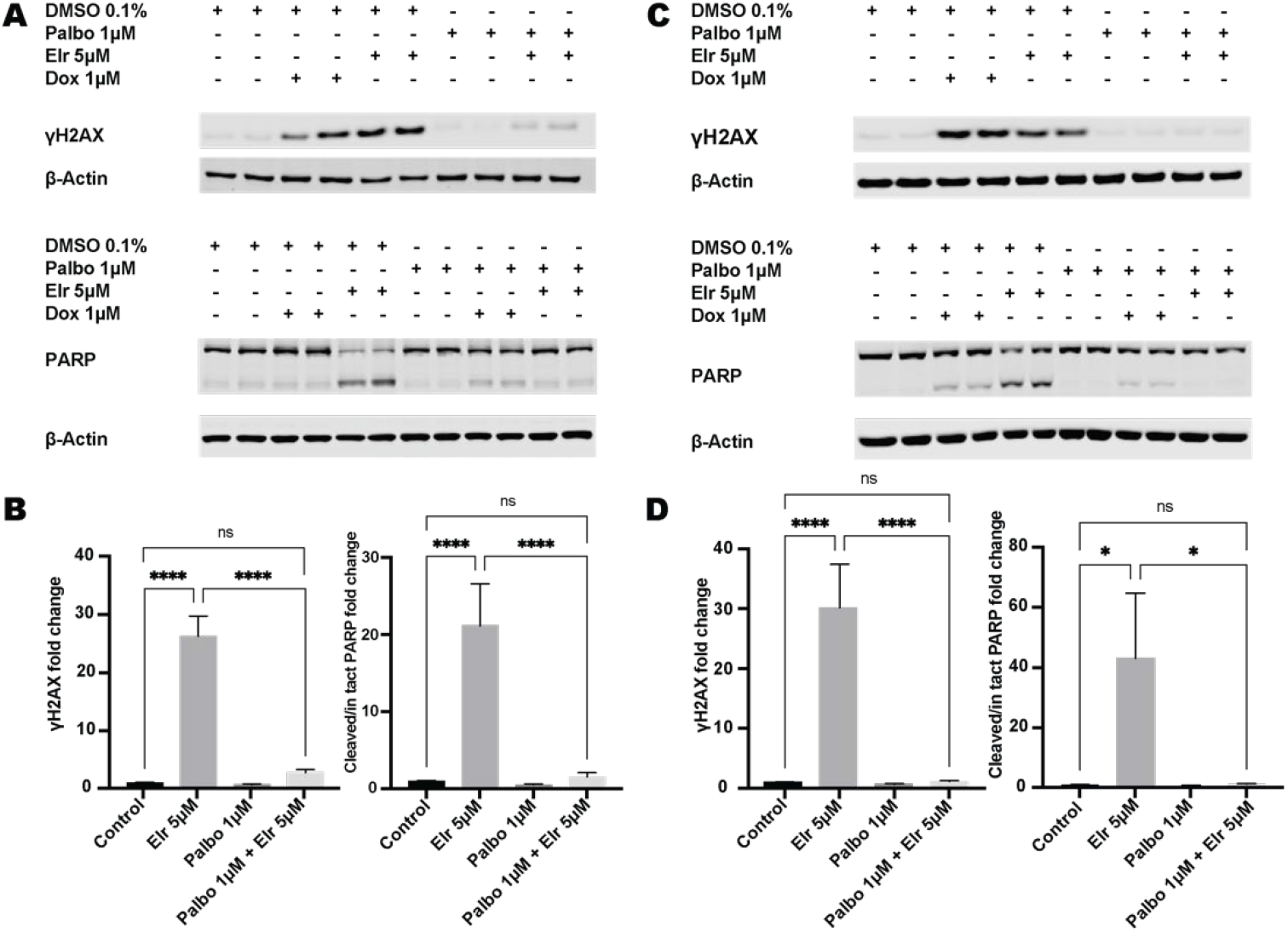
Elraglusib-induced genotoxicity and cytotoxicity are abolished by cell cycle arrest (A-D). Karpas-299 (A, B) and HH cells (C, D) were arrested in palbociclib (Palbo) (or treated with DMSO control) for 24 hours before the addition of elraglusib (Elr), doxorubicin (Dox) or DMSO for a further 24 hours. Cells were then lysed and analysed by immunoblot. PARP cleavage was used for apoptosis assessment and γH2AX for DNA damage. Quantification was performed by normalising to actin and the experimental control. Blots are representative of 3 independent experiments, and quantification combines 3 independent experiments. Fold change calculated versus control, error bars are SEM, statistical significance by one-way ANOVA p ≤0.05.

### Elraglusib impairs the proliferation of primary human T-cells

Elraglusib produces cytotoxic effects in T-cell lymphoma lines (6, 17). Therefore, we sought to compare if these effects were also observed in primary T cells. Figure 5 shows that the viability of unstimulated primary human T-cells was not affected by elraglusib at concentrations of 0.1, 1 or 5µM, but was reduced by 10µM (Fig. 5A). In contrast, primary human T-cells activated and stimulated by CD3/CD28 co-stimulation were affected by elraglusib in a dose-dependent manner at concentrations at or above 1µM (Fig. 5B). Tubulin fractionation immunoblots performed on proliferating primary human T-cells showed that elraglusib led to an increase in soluble tubulin fraction and a reduction in the insoluble tubulin fraction at similar concentrations that induced cytotoxicity (≥1µM) (Fig. 5C, D). CT99021 5µM had no significant effect on soluble or insoluble tubulin fractions in these cells. We conclude that elraglusib affects primary human T cells at similar concentrations to lymphoma lines. These effects require T cell activation, consistent with proliferation being needed to drive cells through mitosis and induce mitotic slippage.

**Figure 5.**
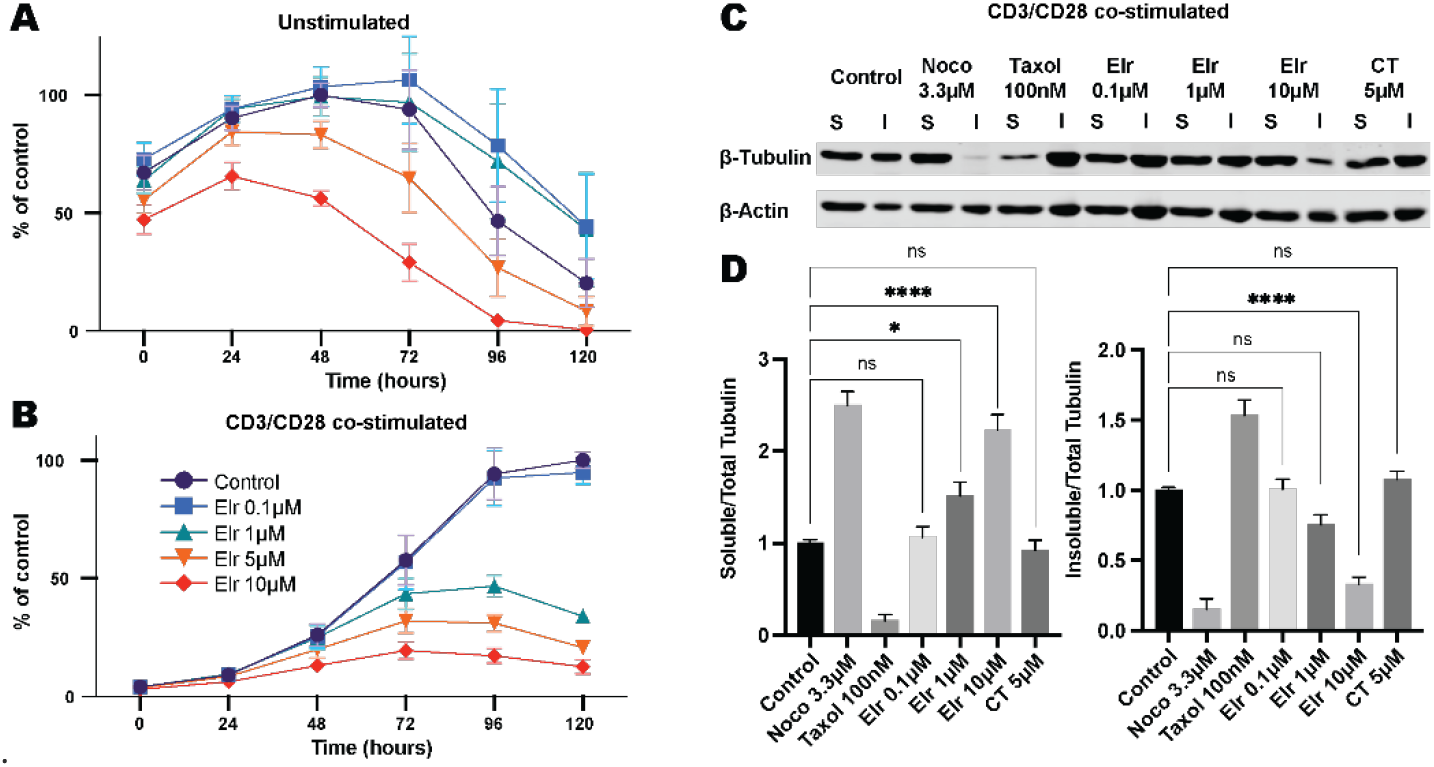
Elraglusib impairs proliferation and reduces cell viability in stimulated primary human T-cells but does not affect unstimulated cells (A, B). Primary human T-cells were isolated from PBMCs from healthy volunteers. Unstimulated controls (A) remained in complete media, stimulated cells (B) were treated with a CD3/CD28-co-stimulation kit. Cells were treated with DMSO control or elraglusib (Elr). Cell viability was assayed using Promega RealTime-Glo assay and the plate read daily for 120 hours. Data combines 3 independent experiments with T-cells from 3 individual volunteers. Data normalized to the experimental control peak luminescence. Elraglusib increases the soluble tubulin fraction and decreases the insoluble tubulin fraction in stimulated primary human T-cells **(C, D)**. Proliferating stimulated primary human T-cells were treated for 5 hours and assayed for soluble and insoluble tubulin as described above. Blots are representative of 3 independent experiments, and quantification combines 3 independent experiments. Fold change calculated versus control, error bars are SEM, statistical significance by one-way ANOVA p ≤0.05.

## Discussion

Elraglusib was developed as a targeted GSK3β inhibitor, and its anti-cancer effects have been variously attributed to GSK3 inhibition within cancer cells, resulting in abnormal regulation of NFkB, impaired DNA-damage responses or myc downregulation, and to immunomodulatory effects within NK and T-cells (7, 13, 24, 25). We, and others, have reported that the drug inhibits both the α and β paralogs of GSK3, along with several other kinases in cell-free assays (10, 17). We also showed that other validated small molecule GSK3α/β inhibitors do not share the anti-lymphoma properties of elraglusib and that *GSK3A/B* shRNA knockdown did not affect the elraglusib IC_50_ (17, 18). Preprint data from Cosgun *et al* also reports broad cytotoxic effects of elraglusib in the absence of β-catenin or myc stabilisation, suggesting it occurs without significant GSK3 inhibition (17, 26). Together, these findings question the requirement for GSK3 inhibition in the mechanism of action of elraglusib

We report here that elraglusib causes a mitotic arrest in various different cancer lines, consistent with the G2 or M arrest observed previously in lymphoma, renal, and bladder cancer lines (6, 9, 10). Importantly, this arrest is associated with direct microtubule depolymerisation which prevents chromosomal alignment leading to mitotic slippage, DNA damage and apoptosis. The process of mitotic slippage is likely to explain the significant increase in non-mitotic H3-pS10 negative 4N cells that appear in the cell cycle analysis of elraglusib-treated cells (Fig. 1). It is also likely to be crucial for the resulting DNA damage and apoptosis because the genotoxic effects of elraglusib are abolished by prior inhibition of CDK4/6 to prevent cell cycle progression.

Similar effects are commonly observed with other anti-cancer agents that block microtubule polymerisation, such as the alkaloids vincristine and vinblastine, and nocodazole (27-29). A recent genome-wide CRISPR/Cas9 knockout screen identified that aurora kinase A knockout or inhibition sensitizes pancreatic cancer cell lines to elraglusib (30). Interestingly, Aurora kinase A inhibition has previously been shown to synergize with vincristine, a recognized microtubule destabilizer, suggesting a shared mechanism of action (31).

The ability of elraglusib to induce mitotic slippage, DNA damage and apoptosis, suggests that it acts similarly to a range of other anti-mitotic drugs that delay mitosis by activating the SAC. These include microtubule stabilizers, such as paclitaxel (28, 32), inhibitors of microtubule motors, such as Eg5 (33, 34), or inhibitors of mitotic kinases, such as PLK1 (35). These drugs all cause side effects due to off-target effects on proliferating healthy cells, especially within the hematopoietic precursor cell compartment, which limits their clinical tolerability. So, could elraglusib offer benefits over commonly used anti-mitotic drugs?

Recently presented trial data from pancreatic cancer patients treated with elraglusib shows a correlation between treatment-related neutropenia and survival, suggesting that off-target effects on healthy cells are probably inevitable to achieve good treatment responses (36). Elraglusib has been reported to specifically target lymphoma cells without affecting the viability of unstimulated normal B- or T-lymphocytes, which could in theory help lymphocytes to mount an anti-tumour immune response (6). However, our data suggests that this difference is due to a lower rate of proliferation in unstimulated lymphocytes, because stimulating the proliferation of primary T lymphocytes enhances elraglusib toxicity (Fig. 5) and preventing proliferation of lymphoma lines abolished cytotoxicity (Fig. 4). The *in vitro* CD3/CD28 co-stimulation model we utilized to stimulate T lymphocytes is designed to mimic T-cell responses to antigen-presenting cells *in vivo*, suggesting that elraglusib may have immunosuppressive effects in patients (37). However, it is important to note that elraglusib was recently shown to enhance immune cell activation and synergize with anti-PD-L1 therapy in a mouse model of colorectal cancer (24). Whether these effects are due to direct effects on the immune system or enhanced immune engagement due to cytotoxic effects on colorectal cancer cells remains to be determined.

An interesting clinical observation that does appear to separate elraglusib from other microtubule-based drugs, concerns the relative lack of neuronal side effects. Microtubule poisons affect axonal transport leading to chemotherapy-induced peripheral neuropathy, and these are frequent dose-limiting toxicities for both vinca alkaloids and taxanes (38). Therefore, it is noteworthy that neuropathy has not yet emerged as a significant side effect of elraglusib (15). Whether this is due to a unique mechanism of action, early-phase data with small patient cohorts, poorer tubulin engagement, or a dosing effect is unclear currently and requires further evaluation.

Elraglusib is not unique in its ability to inhibit both kinases and tubulin polymerisation. Tivantinib was developed as a MET inhibitor, received FDA orphan status and underwent a phase III clinical trial for the treatment of hepatocellular carcinoma (39). Its mechanism does not rely on MET inhibition but depends upon its ability to act as a microtubule destabilising agent (40, 41). Rigosertib, initially identified as a PLK1 inhibitor, but then proposed to function as a RAS mimetic, has also been found to act as a microtubule destabilizer (42-45). Crystallography confirmed that the compound binds tubulin, and specific β-tubulin mutations induce rigosertib resistance without conferring resistance to vinblastine. Conversely, nocodazole, which is not used clinically, is widely used as a microtubule destabilising compound but also inhibits several kinases, including ABL, c-KIT, BRAF and MEK (46).

Off-target effects frequently contribute to the mechanism of action of cancer drugs in clinical trials, and incorrectly attributing the mechanism may confound patient selection, leading to misinterpretation of results and increasing the risk of trial failure (47). Altering a candidate compound to optimize pharmacodynamic and pharmacokinetic properties is often required as part of lead optimisation, and maintaining target engagement requires knowledge of the relevant target (48). Similarly, the identification of a predictive biomarker, which relies upon detailed biological understanding, significantly increases the chance of clinical trial success and FDA approval (49). A detailed understanding of drug mechanisms is also required to appreciate the therapeutic selective pressures which drive resistance, a common cause of treatment failure (50).

Early-phase trial evidence demonstrates the potential clinical efficacy of elraglusib in pancreatic cancer, melanoma, and ATLL (14-16). Our work suggests that these anti-cancer effects occur via microtubule destabilisation rather than GSK3 inhibition. This has implications for the development of GSK3 expression or activity assays as predictive biomarkers for elraglusib response, the evaluation of potential drug resistance mechanisms, the assessment of anticipated patient toxicities, and the design of synergistic drug combinations in future clinical trials.

## Conclusion

Elraglusib acts as a potent direct microtubule destabilizer both *in vitro* and across cell types, resulting in mitotic arrest, genotoxicity and apoptosis. These effects account for its broad pan-cancer cytotoxicity, which does not rely upon GSK3 inhibition and cannot be replicated by other GSK3 inhibitors. This mechanistic data is important for future compound development, clinical trial design, patient toxicity assessment and predictive biomarker development, which should not focus on GSK3-related mechanisms alone.

## Supporting information

Supplementary Movie 1 - 1um Elraglusib

Supplementary Movie 2 - DMSO control

## Author contributions

SL performed the *in* vitro microtubule assay experiments under the supervision of TT. JTC performed all other experiments under the supervision of ST, CS, and ATS. JTC and ATS wrote the manuscript with input from SL, TT, ST and CS.

## Acknowledgements

We thank the staff at the Dundee Imaging Facility and Immunoassay Biomarker Core Laboratory at the School of Medicine. We also thank Dr Rob Wolthuis, Amsterdam UMC, for kindly providing the U2OS H2B-GFP/mCherry-α-tubulin cells. This work was funded by the Ninewells Cancer Campaign Fraser Fellowship Doctoral Training Programme and the NHS Tayside Charitable Foundation (JC, ST, CS), Wellcome Trust Investigator Award (219418/Z/19/Z) and BBSRC grant (BB/X014517/1) (SL, TT) and Cancer Research UK Programme Foundation Award (C4730/A21229) (ATS).

## Methods

### Compounds and antibodies

Doxorubicin and elraglusib (9-ING-41) were purchased from Cambridge Bioscience, palbociclib from MedChemExpress, LY2090314, nocodazole and paclitaxel (Taxol) from Stratech, MG132 from Sigma Aldrich, DAPI (4,6-diamidino2-phenylindole) from Thermo Fisher, and RO3306 from Tocris. CT99021 was synthesized locally as previously described (51). Compounds were dissolved in dimethyl sulfoxide (DMSO) before use. Flow cytometry antibody: rabbit anti-phospho-histone 3 (Ser10) (Cell Signaling 3465) used at 1:100. Immunoblot antibodies: rabbit anti-β-tubulin (Cell Signaling 2128), rabbit anti-PARP (Cell Signaling 9542), rabbit anti-phospho-histone H2A.X (γH2AX) (Cell Signaling 9718), rabbit anti-β-Catenin (Cell Signaling 9562) were all used at 1:1000. Mouse anti-actin (Sigma Aldrich A3853) was used at 1:2000. IF antibodies: mouse anti-α-Tubulin (Sigma Aldrich T5168) was used at 1:2500. Guinea pig anti-CENPC (MBL PD030) was used at 1:5000. Mouse anti-MAD1 (Merck Millipore MABE867) was used at 1:1000. Secondary antibodies for immunoblot: goat anti-rabbit IRDye 800CW (LI-COR 926-32211), goat anti-mouse Alexa Fluor 680 (Invitrogen A21057) both used at 1:10,000. Secondary antibodies for IF: chicken anti-mouse Alexa Fluor 488 (Invitrogen A21200), donkey anti-rabbit Alexa Fluor 568 (Invitrogen A10042), goat anti-guinea pig Alexa Fluor 647 (Invitrogen A21450) all used at 1:1000.

### Cell Culture

K299 cells were purchased from Public Health England, MCF7 and HH cells were from American Type Culture Collection (ATCC). U2OS H2B-GFP/mCherry-α-tubulin were a gift from Rob Wolthuis and were generated as described previously (52). Suspension cells were cultured in RPMI-1640 (Gibco) media containing glycine supplemented with 10% fetal bovine serum (FBS) (Sigma) and 100 units/ml penicillin, 100µg/ml streptomycin (Gibco). Adherent cells were cultured in Dulbecco’s modified Eagle media (Gibco) supplemented with 10% FBS (Sigma) with 100 units/ml penicillin, 100µg/ml streptomycin (Gibco). All cells were maintained at 37°C in an atmosphere of 5% CO_2_. Cells were periodically screened for mycoplasma infection by PCR.

### Primary human T-cell isolation and proliferation

Primary human T-cells were isolated from peripheral blood mononuclear cells (PBMCs) taken from consenting healthy volunteers. Ethical approval was provided by the University of Dundee School of Medicine and Life Sciences Research Ethics Committee (application number - UOD-SMED-SLS-RPG-2024-24-02). PBMCs were isolated from venous blood drawn into sodium heparin vacutainer tubes (BD) using SepMate tubes (STEMCELL Technologies) and T-cells purified using a MACS (magnetic activated cell sorting) negative-selection Pan-T-cell isolation kit (Miltenyi) as per the manufacturers’ instructions. Briefly, Ficoll-Paque PLUS density gradient media (Cytiva) was pipetted into SepMate tubes before the addition of blood (diluted in PBS (1:1 v/v)). Following centrifugation and PBS washes, the pelleted PBMCs were counted, and the appropriate volume of antibody and microbead cocktails was added prior to MACS. T-cells were then cultured as per other suspension cells (see above).

Rapid-Act T-cell activation kit (human, anti-CD3/CD28) (Cell Signaling) was used to induce T-cell proliferation as per the manufacturer’s instructions. Briefly, T-cells were counted, and the appropriate volume of Rapid-Act CD3/CD28 complex was added. Activated T-cells were either incubated until actively proliferating for tubulin experiments or immediately plated for cell viability assay experiments. RealTime Glo (Promega) cell viability assay was used to compare unstimulated and stimulated T-cells as per the manufacturer’s instructions. Briefly, 30,000 cells were added per well of a white-walled 96-well plate (Thermofisher) in 100µL complete media. RealTime Glo reagents were added (1:1000) and the plate was returned to the incubator for 10 minutes before a baseline luminescence read and then read every 24 hours for 120 hours. Media-only wells were used to subtract background luminescence. 3 individual experiments were performed using T-cells from 3 separate volunteers, each experimental condition was done in technical triplicate. Data was normalized to the control peak luminescence for unstimulated and stimulated groups.

### Flow cytometry

4 million cells/well were treated for 24 hours in 6-well plates and then fixed in ice-cold 70% ethanol for 20 minutes at 4°C. Cells were permeabilized by resuspension in 1ml 0.25% Triton-X100 in PBS on ice for 15 minutes. Following PBS washes, cells were re-suspended in anti-phospho-H3 antibody for 1 hour in the dark, then washed in 1% BSA in PBS and then PBS. Cells were resuspended in 50µL ribonuclease A (100µg/ml) (Thermofisher) in PBS and 300µL propidium iodide (50µg/ml) (Thermofisher). Samples were shielded from light and incubated for 20 minutes at room temperature.

A BD LSRFortessa flow cytometer and FACSDiva software were used to process samples. FSC-A and SSC-A plots were produced, and single-cell populations were identified. 10,000 events were recorded per sample. Negative unstained controls were run alongside experimental samples. FCS files were analysed using open-source Floreada.io software and GraphPad Prism. Cell cycle histograms are representative of 3 independent experiments, quantification combines 3 independent experiments. Fold change calculated versus control, error bars are SEM, statistical significance by one-way ANOVA p ≤0.05.

### Live cell imaging

U2OS H2B-GFP/mCherry-α-tubulin cells were incubated overnight in an Ibidi 8-well chamber slide. Immediately before live-cell imaging the media was replaced with compounds dissolved in complete media (CM). Cells were imaged at 5-minute intervals for 24 hours on a Zeiss Axio Observer using a Plan-apopchromat 20x/0.8 M27 air objective while being maintained at 37°C in an atmosphere of 5% CO_2_. For mitotic duration and mitotic outcome quantifications 10 cells were analysed per field of view with two fields per condition for each experiment. Images were analysed using ImageJ software. Duration and outcome were assessed independently on different randomly selected cells. Mitotic duration was calculated from cell round-up until either two daughter cells were produced or mitotic slippage occurred, if the movie ended prior to either outcome this is indicated on the graph (Fig 2. A). Mitotic outcome was assessed morphologically. Plots combine 2 independent experiments.

### Immunofluorescence microscopy (IF)

MCF7 cells were plated on High-Precision 1.5H 12mm coverslips (Marienfeld) in 24-well plates in 1ml CM. For tubulin experiments, coverslips were treated for 24 hours with compounds in CM.

For microtubule-kinetochore (MK) attachment experiments, coverslips were treated with palbociclib 1µM for 24 hours, washed once with PBS, then every 30 minutes for 6 washes with CM, and allowed to recover for 16 hours. CM was replaced with RO3306 10µM in CM for 2 hours before three quick CM washes. After 15 minutes, when mitotic cells were seen, MG132 10µM in CM was added for 30 minutes. Compounds in CM with MG132 10µM were added for 10 minutes.

Cells were fixed in 4% PFA in PBS for 15 minutes at room temperature. Following three PBS washes, coverslips were blocked in 1ml 3% BSA in PBS with 0.5% Triton-X100 for 30 minutes at room temperature. Coverslips were incubated in primary antibody overnight at 4°C. After three PBS washes, coverslips were incubated in a secondary antibody cocktail with DAPI for 2 hours at room temperature. Following three PBS washes, coverslips were dipped in absolute ethanol, air-dried and mounted with ProLong antifade reagent (Molecular Probes). Coverslips were imaged on a DeltaVision 100x/1.40 NA U Plan S apochromat objective and analysed using ImageJ.

For tubulin IF experiments, 10 mitotic cells were imaged per coverslip, images are representative of 3 independent experiments. For MK experiments, CENPC was used to identify kinetochores, and MAD1 positivity was used to assess kinetochore attachment. Images are representative of 3 independent experiments. Quantification represents MAD1-positive kinetochores from 5 mitotic cells per condition and combines 3 independent experiments.

### Live cell and IF imaging statistical analysis

Figure 2 graphs (A, E) are plotted as violin plots using PlotsOfData (https://huygens.science.uva.nl/PlotsOfData/ (53)). This allows the spread of data to be accurately visualized along with the 95% confidence intervals (thick vertical bars) calculated around the median (thin horizontal lines). to allow statistical comparison between all treatments and timepoints. When the vertical bar of one condition does not overlap with one in another condition the difference between the medians is statistically significant (p<0.05), error bars are SD

### *In vitro* microtubule growth assay

Dynamic microtubules were generated and observed as described previously (54). Briefly, purified tubulin proteins (biotinylated tubulin, rhodamine-labelled tubulin, florescence HiLyte 488 tubulin and unlabelled porcine tubulin) were sourced from Cytoskeleton. To prepare microtubule seeds, a 20μM porcine tubulin mix containing 18% biotinylated tubulin, 12% rhodamine-labelled tubulin, and 70% unlabelled tubulin was incubated with 1mM GMPCPP (Jena BioScience, NU-405S) first on ice and then at 37°C for 30 minutes. To separate the microtubules from non-polymerized tubulins, centrifugation was performed using an Airfuge (Beckman Coulter) for 5 minutes. The microtubules underwent another cycle of depolymerisation and polymerisation with 1mM GMPCPP. Finally, the microtubule seed samples were stored in MRB80 buffer (80mM PIPES, pH 6.8, 1mM MgCl2, and 1mM EGTA) with 10% glycerol.

Coverslips were plasma cleaned using a Carbon Coater (Agar Scientific) and treated with PlusOne Repel-Silane (GE Healthcare) for 10–15 minutes. They were then cleaned further by sonication in methanol and rinsed with ultrapure water. Flow chambers were constructed using the cleaned coverslips and microscopy slides. The chambers were treated with 0.2 mg/ml PLL-PEG-biotin (Surface Solutions) in MRB80 buffer for 5 minutes. After washing with MRB80 buffer, 1% pf127 was flowed through the chamber and incubated for 5 minutes, followed by washing the chamber with MRB80 buffer. Rhodamine-labelled microtubule seeds were incubated with 1mg/ml NeutrAvidin (Thermo Fisher Scientific) for 5 minutes. Microtubule seeds were attached to the coverslips with biotin-NeutrAvidin links. Finally, the chambers were incubated with 1mg/ml κ-casein.

The *in vitro* reaction mixture was prepared in MRB80 buffer, consisting of 12μM tubulin mix (11.5μM unlabelled tubulin and 0.5μM fluorescent HiLyte 488 tubulin), 50mM KCl, 2mM MgCl_2_, 0.1% methylcellulose, 0.5 mg/ml κ-casein, 1mM GTP, 6mM DTT, and oxygen scavenging system (400μg/ml glucose-oxidase, 200μg/ml catalase, 4mM DTT, and 20mM glucose). The mixture was centrifuged for 5 min using Airfuge at room temperature. The supernatant was then mixed with nocodazole and elraglusib of indicated concentrations and added to the flow chamber containing microtubule seeds. The chamber was sealed with vacuum grease and observed at 30°C with TIRF microscopy.

Images of dynamic MTs were acquired by TIRF microscopy with a Nikon Eclipse Ti-E inverted research microscope equipped with diode lasers (405 nm, 488 nm, 561 nm, and 647 nm; Coherent), Acousto-Optic Tunable Filters shutter (Solamere Technology), appropriate filters (Chroma), perfect focus system, the CFI Apochromat TIRF 100x 1.49 N.A. oil objective lens (Nikon), Evolve Delta electron-multiplying charge-coupled device 512x 512 camera (Photometrics) and NIS-Elements AR software.

Images were analysed using ImageJ. Minor imaging drift was corrected using the HyperStackRg plugin and Kymographs were generated in time sequence along a chosen line for an individual MT using the Multi Kymograph plugin on ImageJ. Microtubule growth speeds were calculated by measuring the change in microtubule growth (0.16μm per pixel) over time. Catastrophe frequency (events per minute) was determined by dividing the total number of catastrophe events by the total time microtubules spent growing. Analysis by GraphPad Prism with one-way ANOVA p≤0.05 for statistical significance

### Cell lysates

#### Tubulin

Tubulin fractionation was achieved using a method adapted from Gundersen *et al* (22). Microtubule stabilisation buffer (MSB) was made in dH20 (85mM PIPES, 1mM EGTA, 1mM MgCl_2_, 2M glycerol, 0.5% Triton-X100 and one cOmplete, mini, EDTA-free protease inhibitor cocktail tablet (Merck) per 10ml).

Suspension cells were plated at 4 million cells/well in 6-well plates and treated for 5 hours. Cells were pelleted, washed in PBS and re-suspended in 200µL MSB at room temperature. Lysates were centrifuged at 12,000rpm for 15 minutes. Supernatants were transferred to fresh tubes with 50µL 5x Laemmli’s SDS loading buffer (10% w/v SDS, 250mM Tris-HCl, 50% v/v glycerol, 0.025% (w/v) bromophenol-blue, 5% (v/v) β-mercaptoethanol) and represent the soluble fraction. 250µL of 1x Laemmli’s SDS loading buffer was added to the lysate pellet representing the insoluble fraction.

Adherent cells were plated at 500,000 cells/well in 6-well plates and treated for 5 hours. Following a PBS wash, 200µl of MSB was added to each well. Cells were scraped with a rigid cell lifter, and the lysate was centrifuged as above. As an additional step to increase the yield of insoluble tubulin, 250µl of 1x Laemmli’s SDS loading buffer was added to the insoluble debris remaining in each well of the 6-well plates and incubated at room temperature for 15 minutes. Following centrifugation, the soluble fractions were obtained from supernatant as above. The 1x Laemmli’s buffer from the wells was added to the lysate debris pellet to obtain the insoluble fraction.

Soluble and insoluble lysate samples were heated to 95C for 15 minutes at 750rpm on a thermoshaker and stored at -20°C. 10µL of each sample was loaded per well onto acrylamide gels.

#### PARP/γH2AX

PARP cleavage was used to assess for apoptosis and γH2AX for DNA damage by immunoblot. Cell lysates were prepared for SDS-PAGE as previously described, and protein was quantified by Bradford assay (BioRad) (55).

#### Immunoblots

Tubulin, β-Catenin, and PARP samples were run on 10% SDS-PAGE (sodium dodecyl sulfate-polyacrylamide gel electrophoresis) in tris-glycine running buffer (25mM Tris, 192mM glycine, 0.1% SDS), γH2AX samples were run on 15% SDS-PAGE gels in MES buffer (Thermo Fisher). Proteins were transferred onto nitrocellulose membranes, blocked for 1 hour in 1% BSA in TBS-T (40mM Tris, 150mM NaCl with 0.05% (v/v) Tween 20) and incubated overnight at 4°C in primary antibody. Membranes were washed with TBS-T, incubated in secondary antibody for 1 hour, washed in TBS-T, and imaged. Blots analysis was on the LICOR-Odyssey scanner with quantification on ImageStudio Lite 5.2. Blots are representative of 3 independent experiments; quantification combines 3 independent experiments with normalisation to actin loading control and to experimental control. Analysis by GraphPad Prism with one-way ANOVA p≤0.05 for statistical significance

## References

1. Embi N, Rylatt DB, Cohen P. Glycogen synthase kinase-3 from rabbit skeletal muscle. Separation from cyclic-AMP-dependent protein kinase and phosphorylase kinase. Eur J Biochem. 1980;107(2):519–27.

2. Sutherland C. What Are the bona fide GSK3 Substrates? International journal of Alzheimer’s dizease. 2011;2011:505607.

3. Jope RS, Johnson GV. The glamour and gloom of glycogen synthase kinase-3. Trends Biochem Sci. 2004;29(2):95–102.

4. Frame S, Cohen P, Biondi RM. A common phosphate binding site explains the unique substrate specificity of GSK3 and its inactivation by phosphorylation. Mol Cell. 2001;7(6):1321–7.

5. Quintayo MA, Munro AF, Thomas J, Kunkler IH, Jack W, Kerr GR, et al. GSK3beta and cyclin D1 expression predicts outcome in early breast cancer patients. Breast Cancer Res Treat. 2012;136(1):161–8.

6. Wu X, Stenson M, Abeykoon J, Nowakowski K, Zhang L, Lawson J, et al. Targeting glycogen synthase kinase 3 for therapeutic benefit in lymphoma. Blood. 2019;134(4):363–73.

7. Ding L, Madamsetty VS, Kiers S, Alekhina O, Ugolkov A, Dube J, et al. Glycogen Synthase Kinase-3 Inhibition Sensitizes Pancreatic Cancer Cells to Chemotherapy by Abrogating the TopBP1/ATR-Mediated DNA Damage Response. Clin Cancer Res. 2019;25(21):6452–62.

8. Hilliard TS, Gaisina IN, Muehlbauer AG, Gaisin AM, Gallier F, Burdette JE. Glycogen synthase kinase 3 beta inhibitors induce apoptosis in ovarian cancer cells and inhibit in-vivo tumor growth. Anti-Cancer Drug. 2011;22(10):978–85.

9. Kuroki H, Anraku T, Kazama A, Bilim V, Tasaki M, Schmitt D, et al. 9-ING-41, a small molecule inhibitor of GSK-3beta, potentiates the effects of anticancer therapeutics in bladder cancer. Scientific reports. 2019;9(1):19977.

10. Pal K, Cao Y, Gaisina IN, Bhattacharya S, Dutta SK, Wang EF, et al. Inhibition of GSK-3 Induces Differentiation and Impaired Glucose Metabolism in Renal Cancer. Molecular Cancer Therapeutics. 2014;13(2):285–96.

11. Ugolkov A, Gaisina I, Zhang JS, Billadeau DD, White K, Kozikowski A, et al. GSK-3 inhibition overcomes chemoresistance in human breast cancer. Cancer letters. 2016;380(2):384–92.

12. Ugolkov A, Qiang W, Bondarenko G, Procissi D, Gaisina I, James CD, et al. Combination Treatment with the GSK-3 Inhibitor 9-ING-41 and CCNU Cures Orthotopic Chemoresistant Glioblastoma in Patient-Derived Xenograft Models. Transl Oncol. 2017;10(4):669–78.

13. Karmali R, Chukkapalli V, Gordon LI, Borgia JA, Ugolkov A, Mazar AP, et al. GSK-3beta inhibitor, 9-ING-41, reduces cell viability and halts proliferation of B-cell lymphoma cell lines as a single agent and in combination with novel agents. Oncotarget. 2017;8(70):114924–34.

14. ClinicalTrials.gov [Internet]. Bethesda (MD): National Library of Medicine (US). 20 Sep 2018. Identifier NCT03678883, 9-ING-41 in Patients With Advanced Cancers [updated 7 Dec 2022, accessed 25 Jan 2023]. Available from: https://clinicaltrials.gov/ct2/show/NCT03678883.

15. Carneiro BA, Cavalcante L, Mahalingam D, Saeed A, Safran H, Ma WW, et al. Phase I Study of Elraglusib (9-ING-41), a Glycogen Synthase Kinase-3beta Inhibitor, as Monotherapy or Combined with Chemotherapy in Patients with Advanced Malignancies. Clin Cancer Res. 2024;30(3):522–31.

16. Hsu A, Huntington KE, De Souza A, Zhou L, Olszewski AJ, Makwana NP, et al. Clinical activity of 9-ING-41, a small molecule selective glycogen synthase kinase-3 beta (GSK-3beta) inhibitor, in refractory adult T-Cell leukemia/lymphoma. Cancer Biol Ther. 2022;23(1):417–23.

17. Coats JT, Tauro S, Sutherland C. Elraglusib (formerly 9-ING-41) possesses potent anti-lymphoma properties which cannot be attributed to GSK3 inhibition. Cell Commun Signal. 2023;21(1):131.

18. Coats JT, Tauro S, Sutherland C. Redundancy of Glycogen Synthase Kinase 3 in Lymphoma Cell Viability, Proliferation, and the Cytotoxicity of Elraglusib. Blood. 2023;142(Supplement 1):5802-.

19. Cheng B, Crasta K. Consequences of mitotic slippage for antimicrotubule drug therapy. Endocr Relat Cancer. 2017;24(9):T97–T106.

20. Lara-Gonzalez P, Pines J, Desai A. Spindle assembly checkpoint activation and silencing at kinetochores. Semin Cell Dev Biol. 2021;117:86–98.

21. Brito DA, Yang Z, Rieder CL. Microtubules do not promote mitotic slippage when the spindle assembly checkpoint cannot be satisfied. J Cell Biol. 2008;182(4):623–9.

22. Gundersen GG, Khawaja S, Bulinski JC. Postpolymerization detyrosination of alpha-tubulin: a mechanism for subcellular differentiation of microtubules. J Cell Biol. 1987;105(1):251–64.

23. Foy R, Lew KX, Saurin AT. The search for CDK4/6 inhibitor biomarkers has been hampered by inappropriate proliferation assays. NPJ Breast Cancer. 2024;10(1):19.

24. Huntington KE, Louie AD, Srinivasan PR, Schorl C, Lu S, Silverberg D, et al. GSK-3 Inhibitor Elraglusib Enhances Tumor-Infiltrating Immune Cell Activation in Tumor Biopsies and Synergizes with Anti-PD-L1 in a Murine Model of Colorectal Cancer. Int J Mol Sci. 2023;24(13).

25. Shaw G, Cavalcante L, Giles FJ, Taylor A. Elraglusib (9-ING-41), a selective small-molecule inhibitor of glycogen synthase kinase-3 beta, reduces expression of immune checkpoint molecules PD-1, TIGIT and LAG-3 and enhances CD8(+) T cell cytolytic killing of melanoma cells. J Hematol Oncol. 2022;15(1):134.

26. Cosgun KN, Jumaa H, Robinson ME, Kistner KM, Xu L, Xiao G, et al. Targeted engagement of β-catenin-Ikaros complexes in refractory B-cell malignancies. bioRxiv. 2023:2023.03.13.532152.

27. Jordan MA, Thrower D, Wilson L. Mechanism of inhibition of cell proliferation by Vinca alkaloids. Cancer Res. 1991;51(8):2212–22.

28. Schiff PB, Horwitz SB. Taxol stabilizes microtubules in mouse fibroblast cells. Proc Natl Acad Sci U S A. 1980;77(3):1561–5.

29. Xu K, Schwarz P, Ludueña R. Interaction of Nocodazole With Tubulin Isotypes. Drug Development Research. 2002;55:91–6.

30. Ding L, Maeder E, Zhang C, Weiskittel T, Schmitt D, Mazar A, et al. Abstract 4658: Genome wide CRISPR/Cas9 library screening identifies aurora kinase A as a regulator of elraglusib sensitivity in pancreatic cancer. Cancer Research. 2024;84:4658-.

31. Lentini L, Amato A, Schillaci T, Insalaco L, Di Leonardo A. Aurora-A transcriptional silencing and vincristine treatment show a synergistic effect in human tumor cells. Oncol Res. 2008;17(3):115–25.

32. Sudo T, Nitta M, Saya H, Ueno NT. Dependence of paclitaxel sensitivity on a functional spindle assembly checkpoint. Cancer Res. 2004;64(7):2502–8.

33. Mayer TU, Kapoor TM, Haggarty SJ, King RW, Schreiber SL, Mitchison TJ. Small molecule inhibitor of mitotic spindle bipolarity identified in a phenotype-based screen. Science. 1999;286(5441):971–4.

34. DeBonis S, Skoufias DA, Lebeau L, Lopez R, Robin G, Margolis RL, et al. In vitro screening for inhibitors of the human mitotic kinesin Eg5 with antimitotic and antitumor activities. Mol Cancer Ther. 2004;3(9):1079–90.

35. Lenart P, Petronczki M, Steegmaier M, Di Fiore B, Lipp JJ, Hoffmann M, et al. The small-molecule inhibitor BI 2536 reveals novel insights into mitotic roles of polo-like kinase 1. Current biology : CB. 2007;17(4):304–15.

36. Ugolkov A, Koukol A, Kellinger C, Smith SL, Gallipoli S, Gagnon S, et al. Correlation of therapy-induced neutropenia with survival in patients with metastatic pancreatic cancer treated with GSK-3 inhibitor elraglusib (9-ING-41) in combination with gemcitabine/nab-paclitaxel in the 1801 phase 2 study. Journal of Clinical Oncology. 2024;42(16_suppl):e16311–e.

37. Trickett A, Kwan YL. T cell stimulation and expansion using anti-CD3/CD28 beads. J Immunol Methods. 2003;275(1-2):251–5.

38. Nicolini G, Monfrini M, Scuteri A. Axonal Transport Impairment in Chemotherapy-Induced Peripheral Neuropathy. Toxics. 2015;3(3):322–41.

39. Rimassa L, Assenat E, Peck-Radosavljevic M, Pracht M, Zagonel V, Mathurin P, et al. Tivantinib for second-line treatment of MET-high, advanced hepatocellular carcinoma (METIV-HCC): a final analysis of a phase 3, randomized, placebo-controlled study. Lancet Oncol. 2018;19(5):682–93.

40. Xiang Q, Zhen Z, Deng DY, Wang J, Chen Y, Li J, et al. Tivantinib induces G2/M arrest and apoptosis by disrupting tubulin polymerization in hepatocellular carcinoma. J Exp Clin Cancer Res. 2015;34:118.

41. Aoyama A, Katayama R, Oh-Hara T, Sato S, Okuno Y, Fujita N. Tivantinib (ARQ 197) exhibits antitumor activity by directly interacting with tubulin and overcomes ABC transporter-mediated drug resistance. Mol Cancer Ther. 2014;13(12):2978–90.

42. Jost M, Chen Y, Gilbert LA, Horlbeck MA, Krenning L, Menchon G, et al. Combined CRISPRi/a-Based Chemical Genetic Screens Reveal that Rigosertib Is a Microtubule-Destabilizing Agent. Mol Cell. 2017;68(1):210–23 e6.

43. Jost M, Chen Y, Gilbert LA, Horlbeck MA, Krenning L, Menchon G, et al. Pharmaceutical-Grade Rigosertib Is a Microtubule-Destabilizing Agent. Mol Cell. 2020;79(1):191–8 e3.

44. Gumireddy K, Reddy MV, Cosenza SC, Boominathan R, Baker SJ, Papathi N, et al. ON01910, a non-ATP-competitive small molecule inhibitor of Plk1, is a potent anticancer agent. Cancer Cell. 2005;7(3):275–86.

45. Athuluri-Divakar SK, Vasquez-Del Carpio R, Dutta K, Baker SJ, Cosenza SC, Basu I, et al. A Small Molecule RAS-Mimetic Disrupts RAS Association with Effector Proteins to Block Signaling. Cell. 2016;165(3):643–55.

46. Park H, Hong S, Hong S. Nocodazole is a high-affinity ligand for the cancer-related kinases ABL, c-KIT, BRAF, and MEK. ChemMedChem. 2012;7(1):53–6.

47. Lin A, Giuliano CJ, Palladino A, John KM, Abramowicz C, Yuan ML, et al. Off-target toxicity is a common mechanism of action of cancer drugs undergoing clinical trials. Sci Transl Med. 2019;11(509).

48. Hughes JP, Rees S, Kalindjian SB, Philpott KL. Principles of early drug discovery. Br J Pharmacol. 2011;162(6):1239–49.

49. Wong CH, Siah KW, Lo AW. Estimation of clinical trial success rates and related parameters. Biostatistics. 2019;20(2):273–86.

50. Vasan N, Baselga J, Hyman DM. A view on drug resistance in cancer. Nature. 2019;575(7782):299–309.

51. Finlay D, Patel S, Dickson LM, Shpiro N, Marquez R, Rhodes CJ, et al. Glycogen synthase kinase-3 regulates IGFBP-1 gene transcription through the thymine-rich insulin response element. BMC Mol Biol. 2004;5:15.

52. Voets E, Marsman J, Demmers J, Beijersbergen R, Wolthuis R. The lethal response to Cdk1 inhibition depends on sister chromatid alignment errors generated by KIF4 and isoform 1 of PRC1. Scientific reports. 2015;5:14798.

53. Postma M, Goedhart J. PlotsOfData-A web app for visualizing data together with their summaries. PLoS Biol. 2019;17(3):e3000202.

54. Doodhi H, Kasciukovic T, Clayton L, Tanaka TU. Aurora B switches relative strength of kinetochore-microtubule attachment modes for error correction. J Cell Biol. 2021;220(6).

55. Rena G, Guo S, Cichy SC, Unterman TG, Cohen P. Phosphorylation of the transcription factor forkhead family member FKHR by protein kinase B. J Biol Chem. 1999;274(24):17179–83.

